# Genomic epidemiology of third-generation cephalosporin-resistant *Escherichia coli* from Argentinian pig and dairy farms reveals animal-specific patterns of co-resistance and resistance mechanisms

**DOI:** 10.1101/2023.06.15.545115

**Authors:** Oliver Mounsey, Laura Marchetti, Julián Parada, Laura V. Alarcón, Florencia Aliverti, Matthew B. Avison, Carlos S. Ayala, Cristina Ballesteros, Caroline M. Best, Judy Bettridge, Andrea Buchamer, Daniel Buldain, Alicia Carranza, Maite Cortiisgro, David Demeritt, Maria Paula Escobar, Lihuel Gortari Castillo, María Jaureguiberry, Mariana F. Lucas, L. Vanina Madoz, María José Marconi, Nicolás Moiso, Hernán D. Nievas, Marco A. Ramirez Montes De Oca, Carlos Reding, Kristen K. Reyher, Lucy Vass, Sara Williams, José Giraudo, R. Luzbel De La Sota, Nora Mestorino, Fabiana A. Moredo, Matías Pellegrino

## Abstract

Control measures are being introduced globally to reduce the prevalence of antibiotic resistant (ABR) bacteria on farms. However, little is known about the current prevalence and molecular ecology of ABR in key opportunistic human pathogens such as *Escherichia coli* on South American farms. Working with 30 dairy cattle farms and 40 pig farms across two provinces in central-eastern Argentina, we report a comprehensive genomic analysis of third-generation cephalosporin resistance (3GC-R) in *E. coli*. 3GC-R isolates were recovered from 34.8% (cattle) and 47.8% (pigs) of samples from faecally contaminated sites. Phylogenetic analysis revealed substantial diversity suggestive of long-term horizontal transmission of 3GC-R mechanisms. Despite this, mechanisms such as CTX-M-15 and CTX-M-2 were detected more often in dairy farms, while CTX-M-8 and CMY-2, and co-carriage of amoxicillin/clavulanate resistance and florfenicol resistance were more commonly detected in pig farms. This suggests different selective pressures of antibiotic use in these two animal types, particularly the balance of fourth-versus third-generation cephalosporin use, and of amoxicillin/clavulanate and florfenicol use. We identified the β-lactamase gene *bla*_ROB_ in 3GC-R *E. coli*, which has previously only been reported in the family *Pasteurellaceae*, including farmed animal pathogens. *bla*_ROB_ was found alongside a novel florfenicol resistance gene – *ydhC* – also mobilised from a pig pathogen as part of a new plasmid-mediated composite transposon, which is already widely disseminated. These data set a baseline from which to measure the effects of interventions aimed at reducing on-farm ABR and provide an opportunity to investigate zoonotic transmission of resistant bacteria in this region.

**Importance:** Little is known about the ecology of critically important antibiotic resistance among opportunistic human pathogens (e.g. *Escherichia coli*) on South American farms. By studying 70 farms in central-eastern Argentina, we identified that third-generation cephalosporin resistance (3GC-R) in *E. coli* was mediated by mechanisms seen more often in certain species (pigs or dairy cattle) and that 3GC-R pig *E. coli* were more likely to be co-resistant to florfenicol and amoxicillin/clavulanate. This suggests that on-farm antibiotic usage is key to selecting the types of *E. coli* present on these farms. 3GC-R *E. coli* were highly phylogenetically variable and we identified the *de novo* mobilisation of the resistance gene *bla*_ROB_, alongside a novel florfenicol resistance gene, from pig pathogens into *E. coli* on a mobile genetic element that was widespread in the study region. Overall, this shows the importance of surveying poorly studied regions for critically important antibiotic resistance which might impact human health.

## Introduction

There are very few published reports of third-generation cephalosporin-resistant (3GC-R) *Escherichia coli* from farmed animals in South America. In Uruguay, a 2016 study reported that only 1% of bovine calves excreted 3GC-R *E. coli*, which was conferred by production of CTX-M-15 (1). Positivity rates in Brazilian cattle were 18% of samples in a 2014 survey (2). In Uruguayan pigs, 72% of samples were positive for 3GC-R *E. coli* in 2016, with CTX-M-8, CTX-M-14, CTX-M-15, SHV-12 and CMY-2 identified as mechanisms (1). In Argentinian pigs, 3GC-R *E. coli* were also common in 2017 (82% of samples were positive), caused by multiple CTX-M variants (3). In contrast, a 2012 survey performed in Brazil reported that only 3% of samples from pigs were positive (4), which suggests there has been a rapid rise in 3GC-R *E. coli* prevalence in pigs in the wider region. In more recent years, CTX-M producers have been found in wild animal reservoirs, e.g., vampire bats in Peru, though at lower prevalence than the 48% of sampled livestock that excreted 3GC-R *E. coli* (5). In Chile, however, only 3% of livestock samples were positive in a 2019 survey of small-scale farming operations with low use of cephalosporins (6). Generally, however, given a paucity of data, and variations in sampling structure as well as methods for selecting resistant bacteria and identification of resistance genes in these previous studies, it is difficult to make general conclusions about the prevalence and mechanisms of 3GC-R excreted by production animals in South America.

There is much interest in the possibility that resistant *E. coli* emerging on farms might colonise human populations through the ingestion of food or interaction with environments contaminated with farm animal faeces. It is possible that such zoonotic transmission could exacerbate the rise of resistant infections in humans. Accordingly, countries commonly include reducing antibiotic use in farming as a central pillar of their published AMR action plans. In 2015, the Argentinian government adopted the concept of “One Health”, as promoted by the World Organization for Animal Health and the World Health Organization, and the Argentine Strategy for the Control of Antimicrobial Resistance was formalized. In 2022, a new Law concerning the Prevention and Control of Antimicrobial Resistance was enacted, which specifically aims to ensure the responsible use of antibiotics and regulate issues related to the sale and use of these drugs, both in human and animal health (https://www.boletinoficial.gob.ar/detalleAviso/primera/270118/20220824).

Our aim in this work was to perform large-scale sampling of pig and dairy cattle farms in two regions in central-eastern Argentina to assess the current prevalence and molecular epidemiology – based on whole genome sequencing – of 3GC-R *E. coli* on farms in this under-sampled region. These data might in future be used to test for zoonotic transmission and to measure the impact of initiatives to reduce antibiotic use on farm-level AMR prevalence and ecology.

## Results

### Animal-specific 3GC-R E. coli sample-level positivity and resistance mechanisms

Samples from faecally contaminated environments, effluent and animal drinking water were collected at two separate visits from 40 pig farms (404 samples) and 30 dairy cattle farms (310 samples) between March 22, 2021 and August 30, 2021. The numbers of farms, numbers of samples collected and the percentage of samples positive for 3GC-R *E. coli* from each type of animal and each region are shown in **Figure S1** and **Table 1**. Overall, 33.9% (n=242) of samples were positive for 3GC-R *E. coli*. After accounting for clustering by farm and sample type, there was no statistically significant difference in positivity among samples taken from pig farms compared to dairy farms (OR 0.41; 95% C.I 0.12-1.46). However, samples collected in the Rio Cuarto region were more likely to be positive than those from La Plata (OR 3.62; 95% CI 1.87-7.02) (**Table S1**). Only eight farms (two pig and five dairy farms in La Plata, and one dairy farm in Rio Cuarto) were negative for 3GC-R *E. coli* across all samples collected.

**Table 1.** Details of sample-level positivity for 3GC-R *E. coli* among faecal samples from pigs and dairy cattle

|  | **N of**<br>**Farms** | **N of**<br>**Samples** | **N (%) of 3GC-R *E. coli*-positive**<br>**Samples** | **% of**<br>**Total**<br>**Positive**<br>**Samples** | **N (%) of**<br>**Total 3GC-**<br>**R *E. coli***<br>**Sequenced** |
| --- | --- | --- | --- | --- | --- |
| La Plata Dairy | 22 | 230 | 54 (23.5%) | 22.3% | 34 (21.0%) |
| La Plata Pig | 30 | 300 | 99 (33.0%) | 40.9% | 66 (40.7%) |
| Río Cuarto<br>Dairy | 8 | 80 | 34 (42.5%) | 14.0% | 24 (14.8%) |
| Río Cuarto Pig | 10 | 104 | 55 (52.9%) | 22.7% | 38 (23.5%) |
| TOTAL | 70 | 714 | 242 |  | 162 (48.2%) |

From the 242 positive samples, 162 representative 3GC-R *E. coli* were selected for WGS (**Table 1**), which revealed a wide range of 3GC-R mechanisms, dominated by CTX-M variants (**Fig. 1**). Among the farms with sequenced isolates, four 3GC-R mechanisms demonstrated statistically significant associations with either pig or dairy farms (**Fig. 1**). CMY-2 was found in 25 sequenced 3GC-R isolates from 14 different pig farms but only 3 dairy isolates all from the same farm. (Fisher’s Exact test, p = 0.02, when detection of each 3GC-R mechanism was classified at farm-level). Similarly, CTX-M-8 was found in 40 pig isolates across 22 farms but on only five dairy farms (p=0.01). In contrast, CTX-M-2 was found in only a single pig isolate, but seven dairy farm isolates on five farms (p = 0.02). Most strikingly, CTX-M-15 was detected on six pig farms, but was found in 27 dairy isolates across 14 farms (p < 0.001).

**Figure 1.**
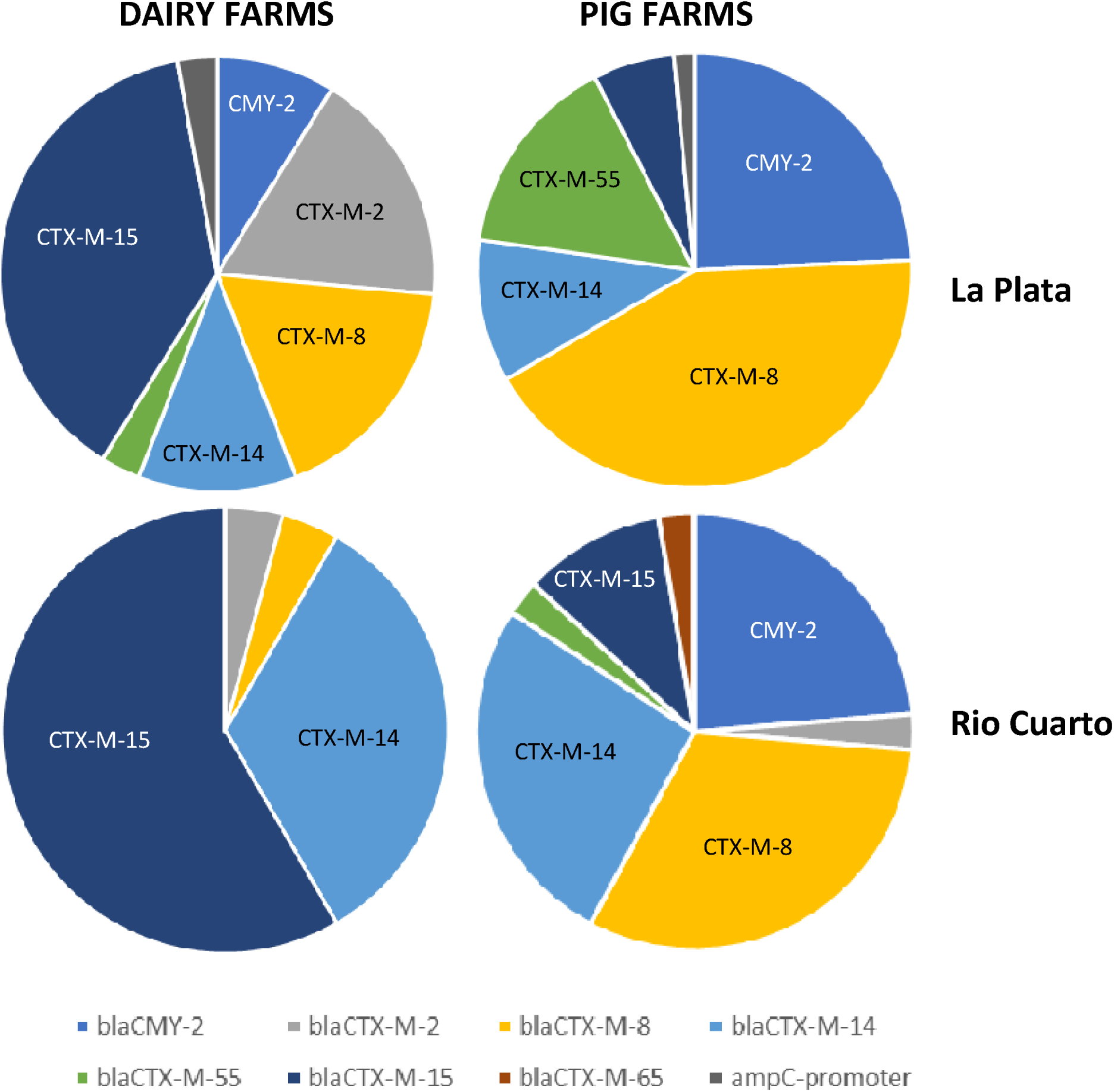
Percentage of 3GC-R *E. coli* isolates from cattle and pigs in Argentina expressing particular 3GC-R mechanisms

### Animal-specific co-resistance phenotypes among 3GC-R E. coli

In terms of other important resistance genes carried among the 162 sequenced 3GC-R *E. coli* isolates (**Table S2**), the florfenicol resistance gene *floR* was more likely to be detected in 3GC-R *E. coli* on pig (65 isolates across 30 farms) than dairy farms (three isolates from three different farms, Fisher’s Exact test, p < 0.001). 3GC-R pig isolates were also far more likely to express amoxicillin/clavulanate resistance. Overall, 54 isolates across 26 pig farms carried one or more amoxicillin/clavulanate resistance mechanism, compared to nine cattle isolates from only 6 farms (p < 0.001). One common mechanism of amoxicillin/clavulanate resistance is production of an AmpC enzyme (7). As previously mentioned, the plasmid mediated AmpC gene *bla*_CMY-2_ was more commonly detected in 3GC-R *E. coli* on pig farms. A second mechanism of amoxicillin/clavulanate resistance is production of an OXA enzyme. Carriage of *bla*_OXA_ genes in 3GC-R *E. coli* was detected on two dairy and six pig farms, which did not suggest a species association (p=0.7) (**Table S2**). A third possible mechanism for amoxicillin/clavulanate resistance is TEM-1 hyper-production. No statistically significant difference in detection of 3GC-R *E. coli* carrying *bla*_TEM-1_-was found from pig farms (70 isolates from 31 farms) compared to dairy farms (28 isolates from 13 farms, p=0.06). However, genetic features associated with TEM-1 hyper-production (7) were observed more commonly (p = 0.003) among *bla*_TEM-1_-positive *E. coli* from pig farms (32 isolates, 20 farms) than from cattle (three isolates, three farms) (**Fig. S2**, **Table S2**). Accordingly, we conclude that amoxicillin/clavulanate resistance was more widespread among 3GC-R *E. coli* from pig farms through higher rates of *bla*_CMY-2_ carriage and higher rates of *bla*_TEM-1_ hyper-expression.

We next considered linkage between specific 3GC-R mechanisms and genes responsible for resistance to other agents. Twenty-two out of 23 *qnr*-positive 3GC-R cattle isolates, representing 16 sequence types (STs) and collected from 10 farms, carried *qnrS1* alongside *bla*_CTX-M-15_ and *bla*_TEM-1_ (**Table S3**). Three out of seven *bla*_CTX-M-15_-positive pig isolates also carried *bla*_TEM-1_ and *qnrS1* (**Table S4**).

Of the remaining five *bla*_CTX-M-15_-positive cattle isolates, three (all ST44) carried *bla*_OXA-1_, *aac(6’)-Ib-cr* and *aac*(*3*)*-IIa* (**Table S3**), with *bla*_OXA-1_ and *aac(6’)-Ib-cr* carried on the same composite transposon (IS*26*-*aac(6’)-Ib-cr*, *bla*_OXA-1,_ *catB*[truncated]-IS*26*). In pigs, all four remaining *bla*_CTX-M-15_-positive isolates were also ST44 and carried *bla*_OXA-1_; two also carried the IS*26*-*aac(6’)-Ib-cr*, *bla*_OXA-1_, *catB*[truncated]-IS*26* transposon but lacked *aac*(*3*)*-IIa* whilst two encoded *bla*_OXA-1_ as part of a shorter transposon (IS*26*-*bla*_OXA-1,_ *catB*[truncated]-IS*26*) and carried *aac*(*3*)*-IIa*.

The only other linkage was found among pig isolates in that nine out of 11 ST48 pig isolates carried *bla*_CMY-2_, representing 36% of all *bla*_CMY-2_-positive pig isolates, though these were distributed across eight farms and both regions.

### Mobilisation of bla_ROB_ into E. coli

One unexpected finding from this study was identification of *bla*_ROB_ in five 3GC-R *E. coli* isolates (**Tables S3, S4**), the first time that this gene has been reported in the *Enterobacterales*. Three isolates were ST48 from three pig farms in the La Plata region, one was an ST3339 isolate from La Plata pigs, and the fifth isolate was ST86, from a Río Cuarto cattle farm. *bla*_ROB_ was found in isolates producing CTX-M-14, –15, –8, CMY-2, TEM-1 and OXA-9. Analysis of the four *bla*_ROB_-positive 3GC-R *E. coli* isolates from pigs identified, in each case, an approximately 6.5 kb composite element not previously seen on the NCBI database (**Fig. 2**). The element is flanked by two copies of IS*26* and includes two composite transposons.

**Figure 2.**
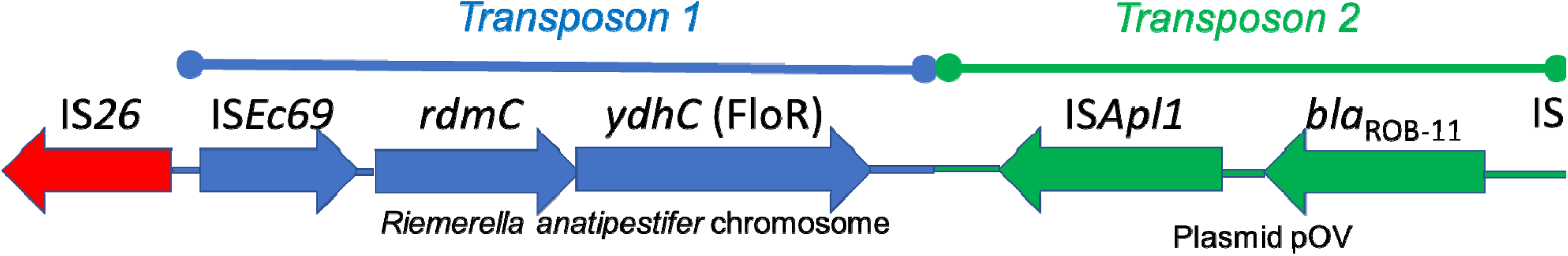
Architecture of a novel mobile genetic element fusing florfenicol and. β-lactam resistance and mobilising *bla*_ROB_ into *E. coli* in Argentina

Transposon 1 includes two adjacent genes, mobilised from the chromosome of the *Flavobacteriales* species *Riemerella anatipestifer* by the IS*1595*-family element, IS*Ec69*. The two genes encoded on transposon 1 are *rdmC*, annotated as encoding an “Alpha/Beta Hydrolase” of unknown function and *ydhC*. Blastn searches revealed that only six *E. coli* NCBI database entries include a gene encoding RdmC with 100% coverage and >95% identity. Four of the six were from pigs or pork in the USA, one was from human faeces in the USA (8) and one was from the USA, but the sample type was not defined. YdhC is 98.5% identical, with 98% coverage, to the archetypal florfenicol resistance transporter FloR (**Fig. S3**) (9) which was identified in all other *bla*_ROB_-negative, FloR-positive 3GC-R *E. coli* isolates found in this study (**Table S2**). The *ydhC* variant of *floR* could not be found outside the chromosome of *R. anatipestifer* using Blastn. Disc susceptibility testing confirmed that all four ROB/YdhC positive 3GC-R pig *E. coli* isolates identified here are florfenicol resistant.

Transposon 2 in this 6.5 kb element (**Fig. 2**) is >99% identical to the classical *bla*_ROB_ transposon from *Pasteurella multocida* plasmid pOV, obtained from a Mexican pig (10), including the IS*Apl1* element that mobilised *bla*_ROB_.

In two out of four *bla*_ROB_-positive pig isolates in this study, WGS data was sufficient to allow prediction of genomic location. In both cases, the 6.5 kb element encoding ROB was located on a plasmid, but in each case the plasmid was different. Based on Blastn, one was highly similar to a variety of IncR *E. coli* plasmids found in samples from pigs from China, Thailand and the UK (e.g. pRHB28-C19_2; accession number CP057369.1) (11). The second was similar to group of IncI1 plasmids, with the closest match being a plasmid from pigs in Portugal (Accession number KY964068.1).

The *bla*_ROB_-positive 3GC-R *E. coli* cattle isolate, which was the only *bla*_ROB_-positive isolate from the Río Cuarto region, did not appear to have the full 6.5 kb ROB-encoding element, lacking the *ydhC* (*floR*) variant, and having an IS*Ec69*-*rdmC* cluster separately from the ROB-encoding transposon. The limitations of short-read sequencing meant we could not confirm plasmid location, but this cattle isolate only carried an IncY plasmid, so the location of *bla*_ROB_ cannot be the same as in the La Plata pig isolates. The mobilised ROB variant was ROB-11, as in the pig isolates.

### Limited evidence of farm-to-farm transmission of 3GC-R E. coli

Across 162 sequenced isolates, 63 previously known STs were identified. Eighteen isolates were from previously unknown STs. These isolates were distributed across 14 new ST designations. The most frequently observed STs were ST10, ST48 – which, as discussed above, was most frequently found in isolates from pigs – ST58 and ST44 (**Table S5**). However, phylogenetic analysis based on core genome alignment revealed that a diverse set of 3GC-R isolates has been recovered in this survey (**Fig. 3**). Forty-seven of the 162 sequenced isolates could be constituted into 21 “clones” where pairs of isolates were separated by <15 core genome SNPs. Of these, only 6 clones were found in samples from two separate farms, and none spanned more than two farms. None of these clones were found in more than one animal species and only one was found in both regions, a pair of ST58 (CTX-M-8) isolates found on pig farms >500 km apart (**Table S6**).

**Figure 3.**
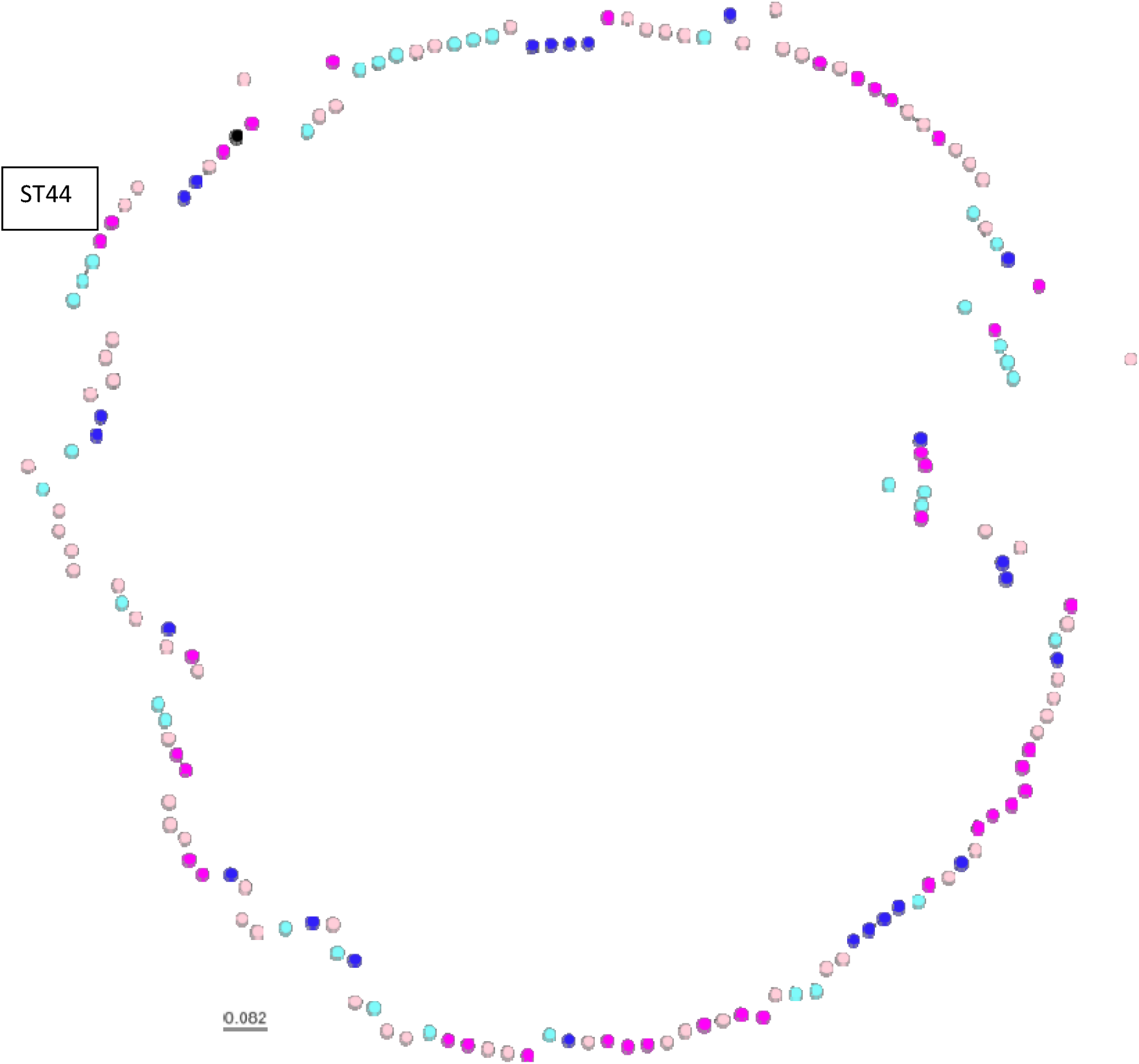
Core genome phylogeny of 3GC-R *E. coli* from Argentinian farms. Pink dots are pig isolates, blue dots are cattle isolates, black is the reference. Dark colours represent isolates from farms near La Plata, light colours represent isolates from farms near Río Cuarto.

Perhaps most importantly, we observed evidence for farm-to-farm sharing of two ST44 (CTX-M-15, IS*26*-*aac(6’)-Ib-cr*, *bla*_OXA-1,_ *catB*[truncated]-IS*26*) clones. One sharing event was identified between Río Cuarto pig farms separated by 20 km; one event, where the plasmid also carried gentamicin resistance, between La Plata dairy farms 4 km apart. These clones were resistant to all first– and second-line antibacterials used for the treatment of bloodstream infections in humans (3GCs, amoxicillin/clavulanate, piperacillin/tazobactam, ciprofloxacin, trimethoprim/sulfamethoxazole, tobramycin, amikacin and – in the La Plata dairy farm clone only – gentamicin). A third ST44 (CTX-M-15, IS*26*-*bla*_OXA-1_, *catB*[truncated]– IS*26*) clone, which also carried gentamicin resistance, was observed in two samples from the same La Plata pig farm. These three ST44 clones differed from each other by >250 SNPs (**Fig. 3**).

## Discussion

There has been little analysis of 3GC-R *E. coli* in samples from production animals in South America, which includes some of the world’s largest and fastest-growing animal farming industries. This is the largest study to date. In pigs and dairy cattle from two regions in central-eastern Argentina, we identified high percentages of sample-level 3GC-R *E. coli* positivity using a method that selects for ESBL– and AmpC-producing *E. coli*. Most isolates recovered, however, produced ESBLs of various CTX-M types. When excluding drinking water samples (where only 1/142 samples was positive for 3GC-R *E. coli*) sample-level 3GC-R *E. coli* positivity for cattle and pigs was 34.8% and 47.8%, respectively. This can be compared with similar studies in other countries, though, as has been discussed in a recent systematic review (12), differences in sampling strategy and selection make comparisons difficult. Using very similar approaches, however, ESBL *E. coli* sample-level positivity was <5% in Canadian cattle (13.14), English dairy farms (15), Australian feedlot beef cattle (16) and Malaysian dairy cattle (17). It is known that animal age is important for finding 3GC-R *E. coli* in cattle (15,18) but the distribution of cattle samples in this study was not substantially biased towards young animals. On the other hand, sample-level 3GC-R *E. coli* positivity rates among cattle reported here were not particularly high compared with the most contemporaneous study from South America – Peru, where positivity was 48% (6). Furthermore, in recent Spanish (19) and Chinese (20) cattle surveillance studies, 3GC-R *E. coli* positivity was 32.9% and 29%, respectively.

In this Argentinian study, 3GC-R *E. coli* positivity seen in pigs was typical of what is seen globally (12). Additionally, we noted that the 3GC-R isolates from pig farms were more likely to express florfenicol and amoxicillin/clavulanate resistance than those from cattle farms. Others have reported more extensive drug resistance in 3GC-R *E. coli* in pigs than cattle when collected in parallel; for example, in South Korea (21,22) and in China (23). In the Argentinian farms studied here, this is likely to reflect the balance of florfenicol and amoxicillin/clavulanate used in pig versus dairy cattle farms, and we aim to provide future epidemiological analysis of 3GC-R *E. coli* presence on these farms following medicines usage and management practice surveys. It was also interesting to note that the type of CTX-M found as a cause of 3GC-R was significantly different between pig and cattle farms in Argentina, with CTX-M-15 (a Group 1 CTX-M) and CTX-M-2 being more common in cattle, and CTX-M-8 being more common in pigs. Indeed, in a parallel sampling study in Italy, CTX-M-15 was also found more commonly to cause 3GC-R in cattle than in pigs (23), though in Canada little difference was seen (25). It may be that any animal-specific imbalance in CTX-M group dominance results from an imbalance between third– and fourth-generation cephalosporin use, since different CTX-M types are known to have different spectra of activity against different cephalosporins (26). A recent study of English dairy cattle also found that Group 1 CTX-M enzymes (in this case CTX-M-32) were the predominant ESBLs seen (27) and showed that the use of fourth-generation cephalosporins was associated with the odds of finding CTX-M-producing *E. coli* in samples from these farms (15). It may be that something similar is occurring in Argentinian dairy farms, but this hypothesis awaits epidemiological evidence. A related observation is that CMY-2 was more commonly found as a cause of 3GC-R in pigs than in cattle, and again this could result from an imbalance in third– and fourth-generation cephalosporin use since CMY-2 does not hydrolyse fourth-generation cephalosporins, as is typical of most AmpC-type enzymes (28).

The other striking finding here is the sheer diversity of 3GC-R *E. coli*, both mechanistically and phylogenetically, suggesting the mobile 3GC-R mechanisms identified have been extensively circulating for many years. Furthermore, 11% of sequenced 3GC-R isolates were from eight previously unknown STs, as might be expected given that this represents a relatively poorly surveyed region of the world. Likely for related reasons, we have identified mobilisation of *bla*_ROB_ into *E. coli* from pigs and, in parallel, mobilisation of a novel florfenicol resistance gene *ydhC*. Despite being detected in both our sampled regions and animal species, ROB has never before been reported in *E. coli* and has exclusively been seen in *Pasterallaceae*, including pathogenic strains from farm animals (29–31) as well as *P. multocida* (32) and *Haemophilus influenzae* (33) from humans, where it is believed to have been mobilised from the chromosomes of bacteria commonly found in pigs (34).

This cements the idea that pig pathogens and commensals are sources of *de novo* resistance gene mobilisation into enteric bacteria with a wider potential for global distribution, as was demonstrated most strikingly with the discovery of the mobile colistin resistance gene *mcr-1* (35). Indeed, in parallel with movement of *bla*_ROB_ into *E. coli*, we also identified *de novo* mobilisation of a novel florfenicol resistance gene *ydhC* from another pig (and avian) pathogen, *R. anatipestifer*, via an IS*Ec69* element, which has previously been associated with composite transposons mobilising the colistin resistance gene, *mcr-2*, which also emerged in pigs (36). This highlights the importance of global resistome surveillance.

The ROB variant encoded in these porcine 3GC-R *E. coli* isolates was ROB-11, which has not formally been characterised before. This variant is one amino acid different from the penicillinase ROB-1 (Thr172Ala) and, whilst this difference is also shared by the reported expanded-spectrum cephalosporinase ROB-2 variant (**Fig. S4**) from *Mannheimia haemolytica*, the role of residue 172 is unclear, and other amino acid changes in ROB-2 are likely more relevant to this phenotype (37). Indeed, ROB-11 was only found in *E. coli* isolates alongside other 3GC-R mechanisms, so there is no reason to believe that this is a novel 3GC-R gene mobilisation. So far, *bla*_ROB_ has not been identified in *E. coli* outside of our study, but this is likely to only be a matter of time.

## Materials and Methods

### Ethical approval and farm sampling

A longitudinal survey of a convenience sample of dairy and pig farms was conducted over the Argentinian autumn and winter seasons of 2021. Farms were located in two regions of central-eastern Argentina, one around La Plata, Buenos Aires province and the other around Río Cuarto, Córdoba province. Requests for farms to enrol in the study were driven by each farm’s proximity to one of two university microbiology laboratories along with previous farmer/veterinarian relationships with the two university veterinary schools undertaking sample collection. During the first visit to each farm, the farmers were informed about the objectives of the project and the number and types of samples to be taken. Then, all farmers gave fully informed consent to participate. Ethical approval was obtained from the Central Bioethics Advisory Committee of the Universidad Nacional de La Plata (13^th^ November 2019).

Among the pig farms selected, 11 were considered small by Argentine standards (< 300 sows in production), 13 medium (301-700 sows) and 16 large (>700 sows). For the dairy farms, at the point of recruitment, 8 were considered small (<200 milking cows), 15 medium (200-500 cows) and 7 large (>500 cows). Each farm was visited once per season, and, at each visit, five samples were collected, returning to the same locations on the farm for each sampling visit. Samples represented a cross-section of sites and animal management stages within farms (**Fig. S1**). Three samples per visit were from faecally-contaminated areas with high densities of animal traffic. For dairy farms, the sites sampled were the calf pen, heifer pen and the waiting area (collecting yard) near the milking parlour. On four large dairy farms with two or three collecting yards, additional samples were taken from these areas.

On pig farms, samples were taken from gestation, weaning (40 days old) and finishing (120 days old) areas. To collect the faecal samples, researchers walked around the areas forming an “X” path, wearing two polyethene absorbent overshoes per collection site which were both stored in a single sterile bag. In addition, for every farm, one sample of effluent outflow and one sample of animal drinking water were collected, in each case using 50 mL sterile polypropylene tubes. One multi-site pig farm had additional effluent and water samples taken from the second site. All samples were kept refrigerated and delivered to the laboratory within 24 h to be processed.

### Sample processing and microbiology

For the overshoe samples, phosphate-buffered saline (PBS) (10 mL/g) was added directly to the pair of used overshoes in their sterile bag and the faecal material manually resuspended. Of this suspension, 1 mL was transferred to a sterile tube, which was centrifuged at 600 rpm for 2 min. The supernatant was diluted 1/200 with PBS and 100 µL were spread onto CHROMagar™ Orientation containing 2 mg/L of cefotaxime to select for 3GC-R *E. coli*. For effluent samples, tubes were left to stand vertically for 30 min, or in some cases centrifuged as above to reduce particulates, then the supernatant was diluted and plated as above. Drinking water samples were centrifuged for 15 min at 3000 rpm and 40 mL of the supernatant were removed; the remaining 10 mL were then used to resuspend any pellet present. Of this suspension, 100 µL were plated as above without dilution.

All plates were incubated for 24 h at 37°C, and two colonies consistent in appearance with 3GC-R *E. coli* (if present) were picked and streaked onto an identical selective agar plate to confirm resistance and to facilitate isolate purification. Purified isolates were then re-checked using Tryptone Bile X-Glucuronide agar plates containing 2 mg/L cefotaxime before PCR and WGS analysis.

### PCR and WGS

Two multiplex PCR assays were used to identify various *bla*_CTX-M_ groups and other β-lactamase genes in 3GC-R isolates as previously (38). Isolates were selected for WGS using these PCR data, with multiple isolates from the same sample only being selected for sequencing if they produced different multiplex PCR profiles. WGS was performed by MicrobesNG as previously (27), and analysed using ResFinder 4.1 (39) and PlasmidFinder 2.1 (40). Hyperexpression of *bla*_TEM-1_ genes was predicted using blastn alignments of upstream sequences to identify known mutations associated with increased promoter activity. Gene copy number was inferred using a bespoke read density analysis, mapping raw sequence reads to a *bla*_TEM-1_ database entry as reference, normalising read density to average chromosomal read density using the MG1655 genome as reference. Details of this approach are presented elsewhere (7).

### Phylogenetics

Sequence alignment and phylogenetic analysis was carried out as described previously (41). SNP distances were determined using SNP-dists (https://github.com/tseemann/snp-dists) and phylogenetic trees were illustrated using Microreact (https://microreact.org/) (42). The reference genome was EC590 – NZ_CP016182.

### Statistical analysis

To estimate any differences in regional or species 3GC-R positivity rates, a binomial mixed-effects model was constructed in R, using the lme4 package (43). Farm of origin and sample type (faecal, effluent or water) were fitted as random effects, and fixed effects were the region, animal species, and visit number, plus an interaction term between species and visit number, based on preliminary findings that detection of 3GC-R *E. coli* tended to be lower in cattle, but not in pigs in visit one compared with visit two.

To test associations between animal species (pig or cattle) and the detection of individual 3GC-R mechanisms, Fisher’s Exact tests were applied, using the data from sequenced isolates. Data were aggregated to give a single presence/absence score for each farm, with subsequent identification of the same mechanism in a different isolate being disregarded, even where sequence types differed.

## Acknowledgements

We are grateful to all farmers and farm workers who facilitated access for sample collection and to Frederico Luna, formerly of Servicio Nacional de Sanidad y Calidad Agroalimentaria, for assistance with project design. Whole genome sequencing was performed by MicrobesNG. We acknowledge the technical assistance of David Griffo, Victorio Fabio Nievas and Walter Darío Nievas.

## Author Contributions

Conceived the Study and Obtained Funding: R.L.S., N.M., J.G., M.P.E., D.D., M.B.A., K.K.R.

Project Management: C.S.A., C.Ba., M.R.M.

Collection of Data: O.M., L.M., J.P., M.P, L.V.A., F.A., A.B., D.B., A.C., M.C.I., L.G.C., D.G., M.J., M.F.L., L.V.M., M.J.M., N.M., H.D.N., supervised by S.W., J.G., M.B.A., K.K.R., R.L.S., F.A.M., N.M.

Cleaning and Analysis of Data: O.M., L.M., J.P., M.P., C.Be., J.B., C.R., L.V., supervised by S.W., R.L.S., N.M., F.A.M., J.G., M.B.A., K.K.R.

Initial Drafting of Manuscript: O.M., M.B.A.

Corrected and Approved Manuscript: All authors

