## Supplementary Information for "Genomic epidemiology of third-generation cephalosporin-resistant *Escherichia coli* from Argentinian pig and dairy farms reveals animal-specific patterns of co-resistance and resistance mechanisms"

**Table S1. Mixed-effects model for 3GC-R *E. coli* positivity in samples from pig and cattle farms across the two study regions**

| | | | | | |
|---|---|---|---|---|---|
| **Random effects** | | | | | |
| | | **Groups** | **Number of groups** | **Variance** | **Std. Dev.** |
| | | Farm | 70 | 0.855 | 0.925 |
| | | Sample type | 3 | | |
| **Fixed effects** | | | | | |
| **Variable** | | **Odds ratio** | **S.E.** | **95% C.I.** | **p** |
| **Intercept** | | 0.20 | 1.45 | 0.01 – 3.33 | 0.26 |
| **Host species** | **Cattle** | Reference category | | | |
| | **Pig** | 0.41 | 0.64 | 0.12 – 1.46 | 0.17 |
| **Region** | **LP** | Reference category | | | |
| | **RC** | 3.62 | 0.34 | 1.87 – 7.01 | <0.001 |
| **Visit number** | **1** | Reference category | | | |
| | **2** | 0.41 | 0.30 | 0.22 – 0.74 | 0.003 |
| **Species: Visit interaction** | | 2.95 | 0.39 | 1.37 - 6.35 | 0.006 |

**Table S2. Other key resistance genes present in sequenced 3GC-R *E. coli* isolates from pig and cattle farms across the two study regions.**

| Source | *N=* | *qnr* | *floR* | *aac(6')-Ib-cr* | *aac(3)/ant(2'')* | TEM-1 (Hyper) | OXA-1 | OXA-9 | ROB-1 |
|---|---|---|---|---|---|---|---|---|---|
| Pig | 104 | 36 | 65 | 4 | 12 | 70 (32) | 4 | 3 | 4 |
| Cattle | 58 | 23 | 3 | 4 | 5 | 28 (3) | 3 | 0 | 1 |
| LP | 100 | 32 | 48 | 6 | 11 | 59 (23) | 5 | 3 | 4 |
| RC | 62 | 27 | 20 | 2 | 6 | 39 (12) | 2 | 0 | 1 |

LP: Farms in La Plata region. RC: Farms in Río Cuarto region. Hyper: TEM-1 predicted to be hyper-produced.

**Table S3. Other key resistance genes carried in 3GC-R sequenced *E. coli* from cattle expressing different mobile 3GC-R mechanisms.**

| 3GC-R Mechanism | *N=* | *STs* | *FARMS* | *qnr* | *floR* | *aac(6')-Ib-cr* | *aac(3)/ant(2'')* | TEM-1 (Hyper) | OXA-1 | OXA-9 | ROB-1 |
|---|---|---|---|---|---|---|---|---|---|---|---|
| CMY-2 | 3 | 2 | 1 (LP) | | | | | | | | |
| CTX-M-2 | 7 | 7 | 4 (LP), 1 (RC) | | 2 | 1 | 2 | 2 (1) | | | |
| CTX-M-8 | 7 | 6 | 4 (LP), 1 (RC) | | | | | 1 (1) | | | |
| CTX-M-14 | 12 | 8 | 3 (LP), 5 (RC) | | | | | 1 (0) | | | |
| CTX-M-15 | 27 | 19 | 8 (LP), 6 (RC) | 23 | 1 | 3 | 3 | 22 (0) | 3 | | 1 |
| CTX-M-55 | 1 | 1 | 1 (LP) | | | | | 1 (0) | | | |

ST: Sequence Type. LP: Farms in La Plata region. RC: Farms in Río Cuarto region. Hyper: TEM-1 predicted to be hyper-produced.

**Table S4. Other key resistance genes carried in 3GC-R sequenced *E. coli* from pigs expressing different mobile 3GC-R mechanisms.**

| 3GC-R Mechanism | *N=* | *STs* | *FARMS* | *qnr* | *floR* | *aac(6')-Ib-cr* | *aac(3)/ ant(2'')* | TEM-1 (Hyper) | OXA-1 | OXA-9 | ROB-1 |
|---|---|---|---|---|---|---|---|---|---|---|---|
| CMY-2 | 25 | 13 | 9 (LP), 5 (RC) | 9 | 17 | 1 | 3 | 16 (7) | | 1 | 1 |
| CTX-M-2 | 1 | 1 | 1 (RC) | 1 | 1 | | 1 | 1 (0) | | | |
| CTX-M-8 | 40 | 24 | 16 (LP), 8 (RC) | 16 | 20 | 1 | 2 | 26 (14) | | 1 | 2 |
| CTX-M-14 | 17 | 12 | 6 (LP, 4 (RC) | 3 | 12 | | 1 | 11 (4) | | 1 | 1 |
| CTX-M-15 | 7 | 4 | 3 (LP), 3 (RC) | 3 | 2 | 2 | 2 | 3 (0) | 4 | | |
| CTX-M-55 | 12 | 8 | 8 (LP), 1 (RC) | 4 | 11 | | 3 | 12 (5) | | | |
| CTX-M-65 | 1 | 1 | 1 (RC) | | 1 | | | 1 (1) | | | |

ST: Sequence Type. LP: Farms in La Plata region. RC: Farms in Río Cuarto region. Hyper: TEM-1 predicted to be hyper-produced.

**Table S5. Phylogenetic groups occurring >5 times among 162 sequenced 3GC-R *E. coli* isolates from pigs and cattle across the two study regions.**

| | ST10 | ST44 | ST48 | ST58 | ST101 | ST1196 | total |
|---|---|---|---|---|---|---|---|
| LP | 15 | 5 | 7 | 4 | 2 | 3 | 76 |
| RC | 2 | 2 | 5 | 10 | 4 | 3 | 55 |
| Cattle | 6 | 3 | 1 | 6 | 1 | 1 | 45 |
| Pig | 11 | 4 | 11 | 8 | 5 | 5 | 86 |

ST: Sequence Type. LP: Farms in La Plata region. RC: Farms in Río Cuarto region.

**Table S6. Clones where sequenced 3GC-R *E. coli* isolates differed by <15 core genome SNPs**

ST 13116 – all same farm: isolates 056954, 056953 and 56912

ST 758 – all same farm: 056975, 056946 and 56809

ST 13108 – all same farm: 056947 and 56797

ST 58 (CTX-M-14) – two dairy farms in RC, 18 km apart: 056950, 056976, 56799 and 56843

ST 2144 – all same farm: 056958 and 56710

ST10 – all same farm: 056961 and 56714

ST7973 (CTX-M-14) – two pig farms in LP: 056965 and 56711

ST1252 – all same farm: 056978 and 56776

ST13104 – all same farm: 56705 and 56706

ST44 (CTX-M-15, IS*26*-*bla*$_{OXA-1}$, *catB*[truncated]-IS*26*, *aac(3)-IIa*) – all same LP pig farm: 56715 and 56717

ST155 – all same farm: 56716 and 56928

ST48 – all same farm: 56718 and 56894

ST29 – all same farm: 56725 and 56726

ST48 – all same farm: 56727 and 56728

ST58 (CTX-M-8) – two pig farms, one in LP, one in RC >500 km apart: 56729 and 56934

ST44 (CTX-M-15, IS*26*-*aac(6')-Ib-cr*, *bla*$_{OXA-1}$, *catB*[truncated]-IS*26*) – two pig farms in RC, 20 km apart: 56732 and 56858

ST1196 – all same farm: 56738 and 56941

ST5229 – all same farm: 56744 and 56846

ST5265 (CTX-M-15) – two dairy farms in RC, 55 km apart: 56792 and 56802

ST10 – all same farm: 56821 and 56822

ST44 (CTX-M-15, IS*26*-*aac(6')-Ib-cr*, *bla*$_{OXA-1}$, *catB*[truncated]-IS*26*, *aac(3)-IIa*) – two dairy farms in LP, 4 km apart: 56841, 56922 and 56924

LP: Farms in La Plata region. RC: Farms in Río Cuarto region.

**Figure S1. Flow diagram representing sample types and numbers collected**

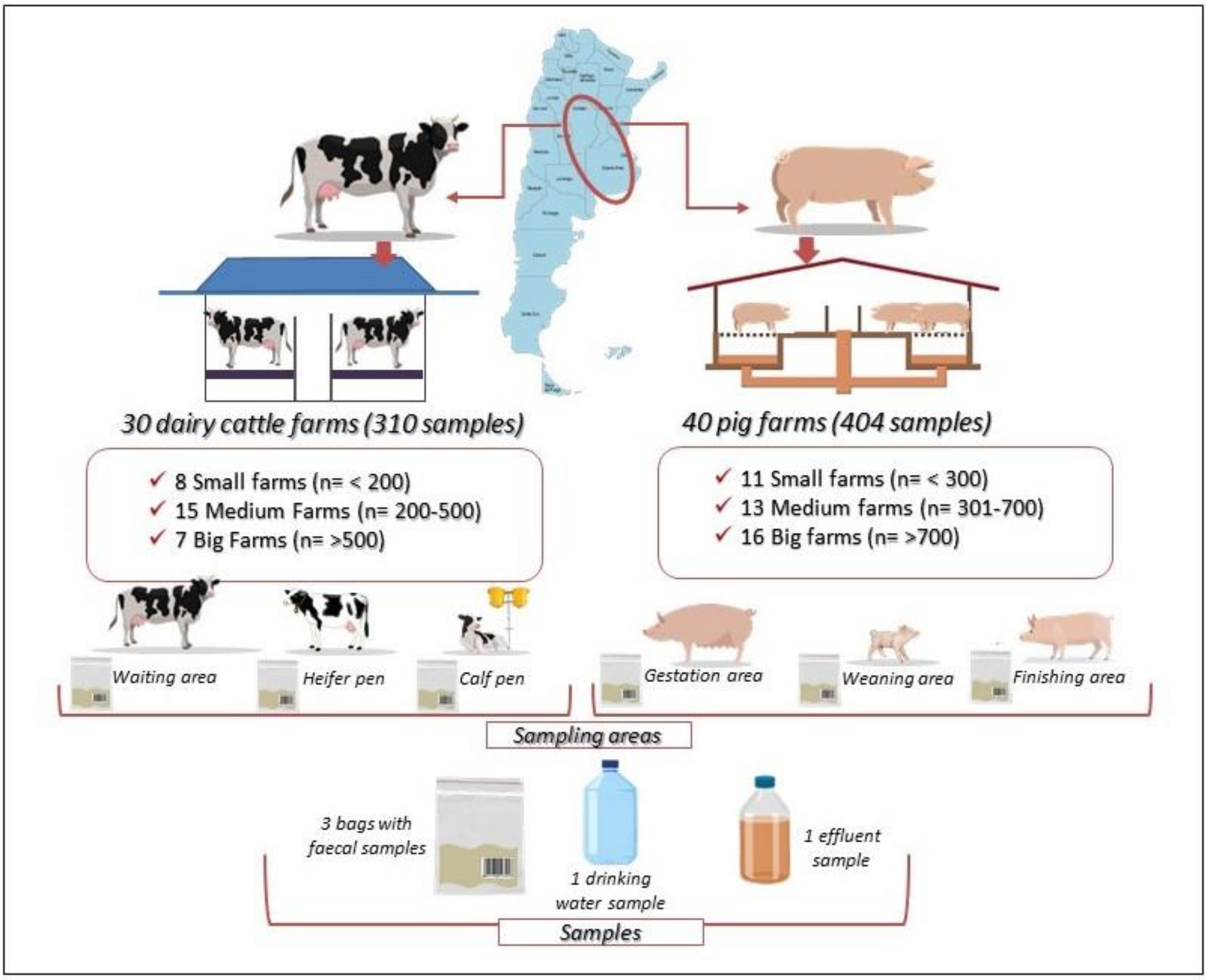

**Figure S2. Analysis of *bla*$_{TEM}$ genes to predict hyper-expression**

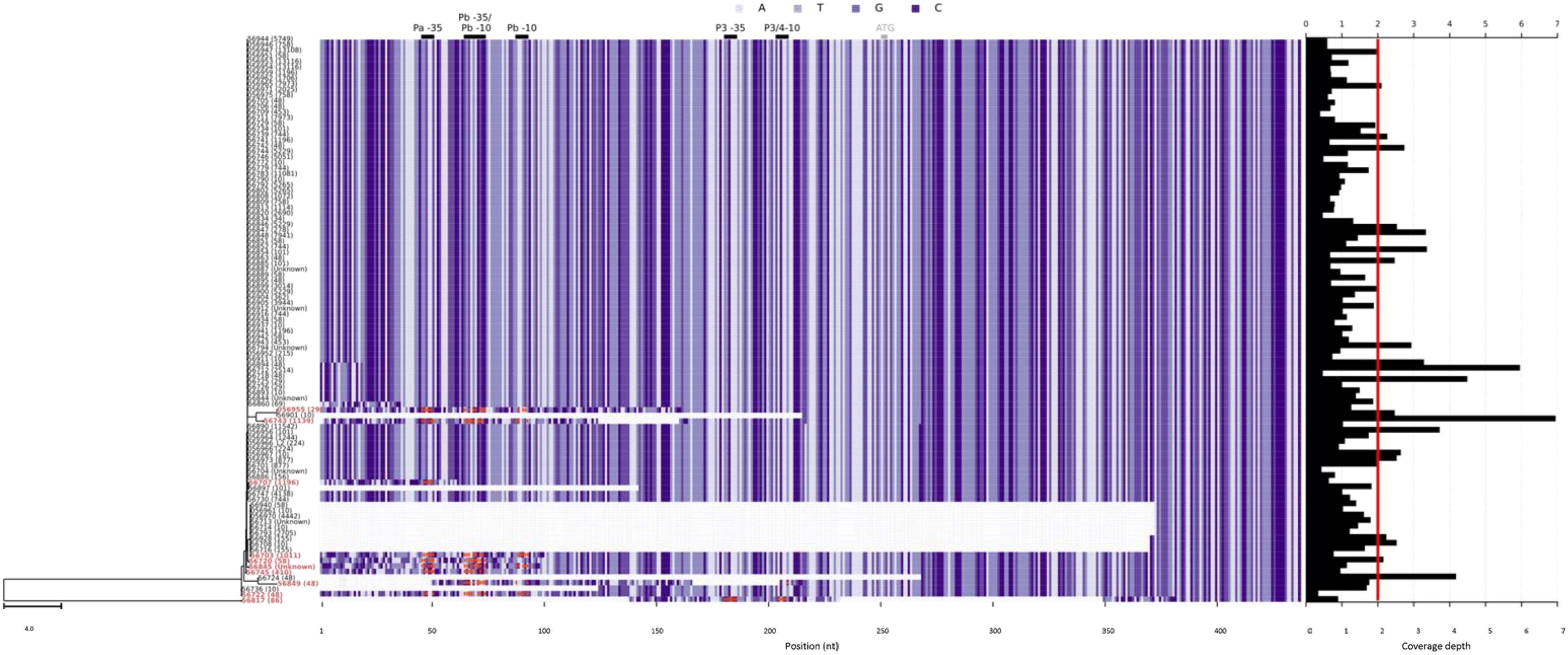

**Figure S3. CLUSTAL alignment of YdhC with the classical florfenicol resistance protein.**

```
YdhC      MTDKQKQRPAWAYTLPAALLLMAPFDILASLAMDIYLPVVPAMPGILNTTPAMIQLTLSL  60
FloR      ---MTTTRPAWAYTLPAALLLMAPFDILASLAMDIYLPVVPAMPGILNTTPAMIQLTLSL  57
               . *****************************************************

YdhC      YMVMLGVGQVIFGPLSDRIGRRPILLAGATAFVIASLGAAWSSTAPAFVAFRLLQAVGAS  120
FloR      YMVMLGVGQVIFGPLSDRIGRRPILLAGATAFVIASLGAAWSSTAPAFVAFRLLQAVGAS  117
          ************************************************************

YdhC      AMLVATFATVRDVYANRPEGVVIYGLFSSILAFVPALGPIAGALIGEFLGWQAIFITLAI  180
FloR      AMLVATFATVRDVYANRPEGVVIYGLFSSMLAFVPALGPIAGALIGEFLGWQAIFITLAI  177
          *****************************:******************************

YdhC      LAMLALLNAGFRWHETRPLDQVKTRRSVLPIFASPAFWVYTVGFSAVMGTYFVFFSTAPR  240
FloR      LAMLALLNAGFRWHETRPLDQVKTRRSVLPIFASPAFWVYTVGFSAGMGTFFVFFSTAPR  237
          ********************************************* ****:*********

YdhC      VLIGQAEYSEIGFSFAFTTVALVMIVTTRFAKSFVARWGIAGCVARGMALLVCGAVLLGI  300
FloR      VLIGQAEYSEIGFSFAFATVALVMIVTTRFAKSFVARWGIAGCVARGMALLVCGAVLLGI  297
          *****************:******************************************

YdhC      GELYGSPSFLTFILPMWVVAVGIVFTVSVTANGALAEFDDIAGSAVAFYFCVQSLIVSIV  360
FloR      GELYGSPSFLTFILPMWVVAVGIVFTVSVTANGALAEFDDIAGSAVAFYFCIQSLIVSIV  357
          ***************************************************:********

YdhC      GTLAVALLNGDTAWPVICYATAMAVLVSLGLVLLRLRGAATEKSPVV   407
FloR      GTLAVTLLNGDTAWPVICYATAMAVLVSLGLALLRSRDAATEKSPVV   404
          *****:*************************.*** *.*********
```

**Figure S4. CLUSTAL alignment of ROB variants.**

```
ROB-2   mlnklkigtlllltltacspnsvhsvksnpqpasapvqqsatqatfqqtlanleqqyqar 60
ROB-1   mlnklkigtlllltltacspnsvhsvtsnpqpasapvqqsatqatfqqtlanleqqyqar 60
ROB-11  mlnklkigtlllltltacspnsvhsvtsnpqpasapvqqsatqatfqqtlanleqqyqar 60
        **************************.*********************************

ROB-2   igvyvwdtetghslsyraderfayastfkallagavlqslpekdlnrtisysqkdlvsys 120
ROB-1   igvyvwdtetghslsyraderfayastfkallagavlqslpekdlnrtisysqkdlvsys 120
ROB-11  igvyvwdtetghslsyraderfayastfkallagavlqslpekdlnrtisysqkdlvsys 120
        ************************************************************

ROB-2   petqkyvgkgmtiaqlceaavrfsdnsatnlllkelggveqyqrilrqlgdnvthanrle 180
ROB-1   petqkyvgkgmtiaqlceaavrfsdnsatnlllkelggveqyqrilrqlgdnvthtnrle 180
ROB-11  petqkyvgkgmtiaqlceaavrfsdnsatnlllkelggveqyqrilrqlgdnvthanrle 180
        *******************************************************:****

ROB-2   pdlnqakpndirdtstpkqmamnlnayllgntltesqktilwnwldnnatgnpliraatp 240
ROB-1   pdlnqakpndirdtstpkqmamnlnayllgntltesqktilwnwldnnatgnpliraatp 240
ROB-11  pdlnqakpndirdtstpkqmamnlnayllgntltesqktilwnwldnnatgnpliraatp 240
        ************************************************************

ROB-2   tswkvydksgagkkygvrndiavvripnrkpivmaimstqfteeakfnnklvedaakqvf 300
ROB-1   tswkvydksgag-kygvrndiavvripnrkpivmaimstqfteeakfnnklvedaakqvf 299
ROB-11  tswkvydksgag-kygvrndiavvripnrkpivmaimstqfteeakfnnklvedaakqvf 299
        ************ ***********************************************

ROB-2   htlqln 306
ROB-1   htlqln 305
ROB-11  htlqln 305
        ******
```
